# A Measure of Open Data: A Metric and Analysis of Reusable Data Practices in Biomedical Data Resources

**DOI:** 10.1101/282830

**Authors:** Seth Carbon, Robin Champieux, Julie McMurry, Lilly Winfree, Letisha R. Wyat, Melissa Haendel

## Abstract

Data is the foundation of science, and there is an increasing focus on how data can be reused and enhanced to drive scientific discoveries. However, most seemingly “open data” do not provide legal permissions for reuse and redistribution. Not being able to integrate and redistribute our collective data resources blocks innovation, and stymies the creation of life-improving diagnostic and drug selection tools. To help the biomedical research and research support communities (e.g. libraries, funders, repositories, etc.) understand and navigate the data licensing landscape, the (Re)usable Data Project (RDP) (http://reusabledata.org) assesses the licensing characteristics of data resources and how licensing behaviors impact reuse. We have created a ruleset to determine the reusability of data resources and have applied it to 56 scientific data resources (i.e. databases) to date. The results show significant reuse and interoperability barriers. Inspired by game-changing projects like Creative Commons, the Wikipedia Foundation, and the Free Software movement, we hope to engage the scientific community in the discussion regarding the legal use and reuse of scientific data, including the balance of openness and how to create sustainable data resources in an increasingly competitive environment.

## INTRODUCTION

In order for biomedical discoveries to be translated into human health improvements, the underlying data must be thoroughly reusable: one should be able to access and recombine data in new ways and make these recombinations available to others. Significant resources and influence have been invested and leveraged to make biomedical data publicly available and scientifically useful [1-3]. Projects such as the NIH NCATS Translator, Data Commons, Illuminating the Druggable Genome, Bgee, and the Monarch Initiative demonstrate that efforts to aggregate and integrate data are seen as a worthwhile undertaking. However, despite these efforts, technical, logistical, descriptive, and legal barriers continue to impede data interoperability and reusability. We are specifically concerned with the ways in which data licensing practices have created widespread legal and financial barriers across the biomedical domain.

While a great number and variety of publicly-funded biomedical data are ostensibly “open”, and some are accessible via aggregated databases, complex licensing issues hinder them from being put to their best use [4-10]. A lack of licensing rigor and standardization forces data users to manually seek, often repeatedly and from multiple data providers, essential reuse and redistributions permissions. Issues include missing licenses, non-standard licenses, and license provisions that are restrictive or incompatible. The legal interpretation of, and compliance with, database license and reuse agreements has become a significant burden and expense for many fields in the scientific community [11]. A complex and lengthy set of legal negotiations is required for a data integration project to legally and freely redistribute all of its relevant data. Ironically, few data resources have the capacity to pursue policy violations and, in our experience, most researchers who restrictively license their data do so because they want to be credited for their work and are unaware of the downstream reuse implications. Thus, it is not uncommon for researchers to ignore license restrictions. This landscape does not benefit data providers, users, or scientific progress.

The (Re)usable Data Project (RDP) originated from discussions on the NCATS Biomedical Data Translator project (https://ncats.nih.gov/translator), which aims to integrate and leverage biomedical information across a vast diversity of sources. While licensing issues influenced many of the reuse barriers we discussed, participants could not agree on licensing standards, illustrating the complexity and confusing state of the data licensing landscape. The RDP was created to systematically describe the current data licensing landscape from the perspective of data aggregation, reuse, and redistribution of publicly funded biological and biomedical data resources. The RDP’s rubric for evaluating data reusability and re-distributability includes a set of criteria and a scoring system that categorizes and weighs licensing and database characteristics, for example the findability and type of licensing terms, negotiation requirements, scope, accessibility, as well as use case and user type restrictions. The RDP aimed to develop a scoring system that is intuitive and comprehensive, but also defensible and agnostic to domain and scientific task. It is important to note that we are not lawyers and the RDP does not provide legal advice. We are a group of scientists, engineers, librarians, and specialists that are concerned about the use and reuse of increasingly interconnected, derived, and reprocessed data. We want to make sure that data-driven scientific endeavors can work with one another in meaningful ways without undue legal concerns. We hope the licensing evaluation rubric will help others navigate the legal synthesis and redistribution of public data and enable data providers to choose licensing terms that make it easier for others to use and redistribute their data.

## METHODS

The RDP’s main efforts have been the creation and application of a rubric that defines the licensing characteristics of aggregated data resources and measures how these licensing behaviors impact reuse. This includes the capture of structured metadata that provide a high-level description of a resource, a working view of its licensing, information to reconstruct the decisions behind our evaluations, and additional notes of interest to others wanting to understand the reusability of a resource’s data. It is important to note that the rubric was constructed for the evaluation of and only applied to public resources databases, not to individual’s datasets or contributions.

### LICENSE CATEGORIZATION

In order to facilitate several points of evaluation in the RDP rubric and illustrate shared qualities among related licenses, the RDP uses an internal categorization of licenses and licensing information, organizing them into six reuse-oriented types. While we acknowledge that the licensing landscape is much more complicated than these categories communicate, classifying licenses via these basic terms was conceptually helpful and provided needed efficiency and simplicity during the evaluation process. The six license types are described and examples are provided below.

### Permissive

Permissive licenses permit reuse, transformation, and redistribution, allowing for attribution. Examples include CC BY 4.0, CC0, and public domain declarations.

### Copyleft

Copyleft licenses allow for reuse, transformation, and redistribution. However, new contributions derived from the original data resource must be distributed under the same license. Examples include CC BY SA 4.0 and the GNU GPL 3.0.

### Restrictive

Restrictive licenses provide more permissions compared to data resources wherein all copyrights have been reserved by the provider, but still include terms that may hinder data integration and reuse. Examples include CC BY ND 4.0.

### Private Pool licenses

A “private pool” license is one where the resource requires data users to add their own data to the pool, or limit the accessibility of derivative data to others that have also joined the pool. Conceptually, this is similar to some copyleft licenses, but without the public “open” component.

### Copyright

This category is used both for licensing statements that positively assert a resource provider’s exclusive copyrights, often referred to as “all rights reserved”, and for when a resource makes no statement about the disposition of its data. Under current US copyright law, creators do not have to explicitly register or copymark their creations to claim their exclusive rights [12].

### Unknown

This category captures licensing statements that have conflicting terms, incompatible license references, or are so non-standard or unclear that a data resource’s reuse terms cannot not be reasonably understood or confirmed.

In our view, only data resources within the permissive category facilitate reuse without negotiation, license alignment, or other burdensome tasks. All other categories have issues that hinder reuse.

### CRITERIA

The RDP’s star rubric (http://reusabledata.org/criteria.html) is a five-part criteria that addresses: the findability and type of licensing terms, the scope and completeness of the licensing, the ability to access the data in a reasonable way, restrictions on how the data may be reused, and restrictions on who may reuse the data. Each of the five criteria (labeled A-E) are quantified by up to a 1.0 star value, so data resource evaluations (e.g., scores) can range from 0 to 5 stars. The rubric is quite extensive, with a branching evaluation workflow, multiple rules and decision points within most parts of the criteria, and bypasses for cases where a particular rule may not apply or make sense.

The first three parts of the criteria (labeled A, B, C) refer to mechanical aspects of license discovery and resource access. Here, standard licenses (i.e., licenses that are invoked referentially or by template, like Creative Commons licenses, Open Database License (ODbL), etc.) are preferred since custom language and terms may require negotiations and possible involvement of institutional counsel to clarify and confirm the rights and permissions. The two latter parts of the criteria (D and E) evaluate the reuse aspects of the licensing terms. Part D considers any restrictions on the *kind* of reuse and part E considers any restrictions on *who* can reuse the data. One star is awarded for each part when all types of reuse are permitted and all audiences can reuse the data without negotiation; however, the rubric does make allowances for some restrictive terms if “research” or “non-commercial” reuse contexts are frictionlessly facilitated. Each part of the criteria can be summarized with the following questions:

#### Criteria A

Is the license or terms of use in an easy-to-find location? Is there one, unambiguous license, as opposed to multiple, conflicting versions? Is the license standard?

#### Criteria B

Does the license clearly define the terms of continuing reuse without need for negotiation with the data creators or resource curators? Does the license have a complete scope that covers all of the data and not just a portion?

#### Criteria C

Does the resource provide its data in a reasonable good-faith location, and is there a reasonable and transparent method of accessing that data in bulk?

#### Criteria D

Are all types of reuse (copying, editing, building upon, remixing, distributing) allowable, with or without attribution?

#### Criteria E

Can any type of user group reuse the data?

It is important to acknowledge that the RDP’s rubric emphasizes the reuse and redistribution needs and activities of U.S. based, non-commercial, research use cases. This perspective reflects our own experience as data resource aggregators and primary data producers, and our frustrations navigating terms of use that limit certain communities’ (e.g., clinical researchers) and kinds of reuse (e.g., new tools) [13]. We also found that this specific and practical point of view was helpful in keeping the rubric and its application logically manageable. When this perspective has limited our evaluations, we have captured the fact that other entities may have different results.

### RUBRIC USAGE

To date, we have fully evaluated 56 data resources with the RDP’s star rubric. As the idea for the RDP emerged from an NCATS Biomedical Data Translator meeting, we originally evaluated data resources used by the Translator and the Monarch Initiative, wherein the reuse and free redistribution of publicly available data for disease discovery has been particularly burdensome. We then expanded our scope to evaluate model organism databases (MODs) and data resources that the newly funded NIH Data Commons Pilot Phase will address [14]. We also evaluated several resources that were brought to our attention by the community.

Each data resource received a score from 0 to 5 stars according to the rubric (http://reusabledata.org/criteria.html). Sources were curated directly into the RDP’s Github repository (https://github.com/reusabledata/reusabledata/tree/master/data-sources) as YAML files from a <u>template</u> to help ensure the provenance of statements. The template includes metadata such as source name, description, source type, license type, data access URL, the license issues uncovered during the evaluation, and any commentary about how the license was evaluated. The evaluations were checked by at least two evaluators, and comments on the evaluations were made on GitHub pull requests to allow for transparency and continued conversation. Evaluations then went through a battery of syntactic and consistency checks. When necessary, the evaluated resource was contacted for clarification.

## RESULTS

### RESOURCE EVALUATION SCORES

Complete evaluations can be viewed on the <u>RDP website</u> and the <u>RDP Github repository</u>.

### Overall scoring

Of the 56 data resources we evaluated, 22 (39%) received between 4 and 5 stars, indicating that they met our broadest requirements for being reusable, which allowed for some caveats, only 10 (18%) received 5 stars meeting all parts (A-E) of the criteria. 23 (41%) of the resources received less than 3 stars, which is notable because even data resources for which the provider has reserved all copyrights can receive a score of 3 stars if the data covered by the license is easily accessible. 32 (57%) of the resources had 3 stars or less, meaning that a majority of resources had significant issues with even basic reusability. Overall, average scores by licensing category were: permissive 4.5, restrictive 2.6, copyright 1.4, unknown 0.7, copyleft 3.0, and private pool 1.0.

### Criteria violations

When a resource provided inconsistent or no licensing information, only parts A and C of the rubric were used in the evaluation. While one could assume that some of these resources wished to reserve all of their copyrights when no information was found, the ambiguity and lack of clear intent would require clarification and possibly legal counsel. 11 data resources had such contradictory or missing information; therefore, the summary statistics for parts B, D, and E of the rubric do not include data for these resources (see Figure 1). We have qualified all of the numbers given below to prevent ambiguity.

**Figure 1.**
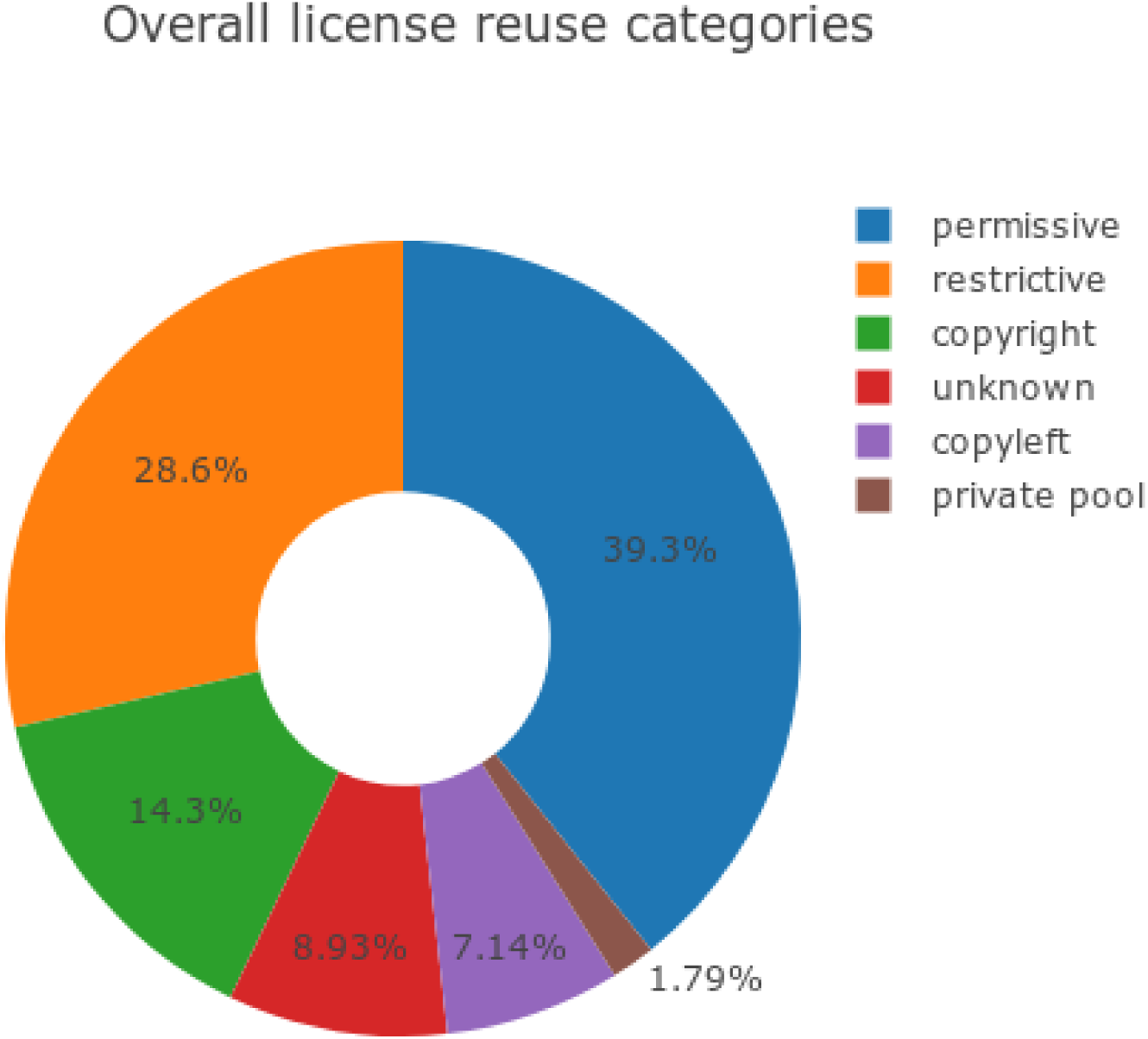
The breakdown of evaluated licenses according to their determined reuse category.

**Figure 2.**
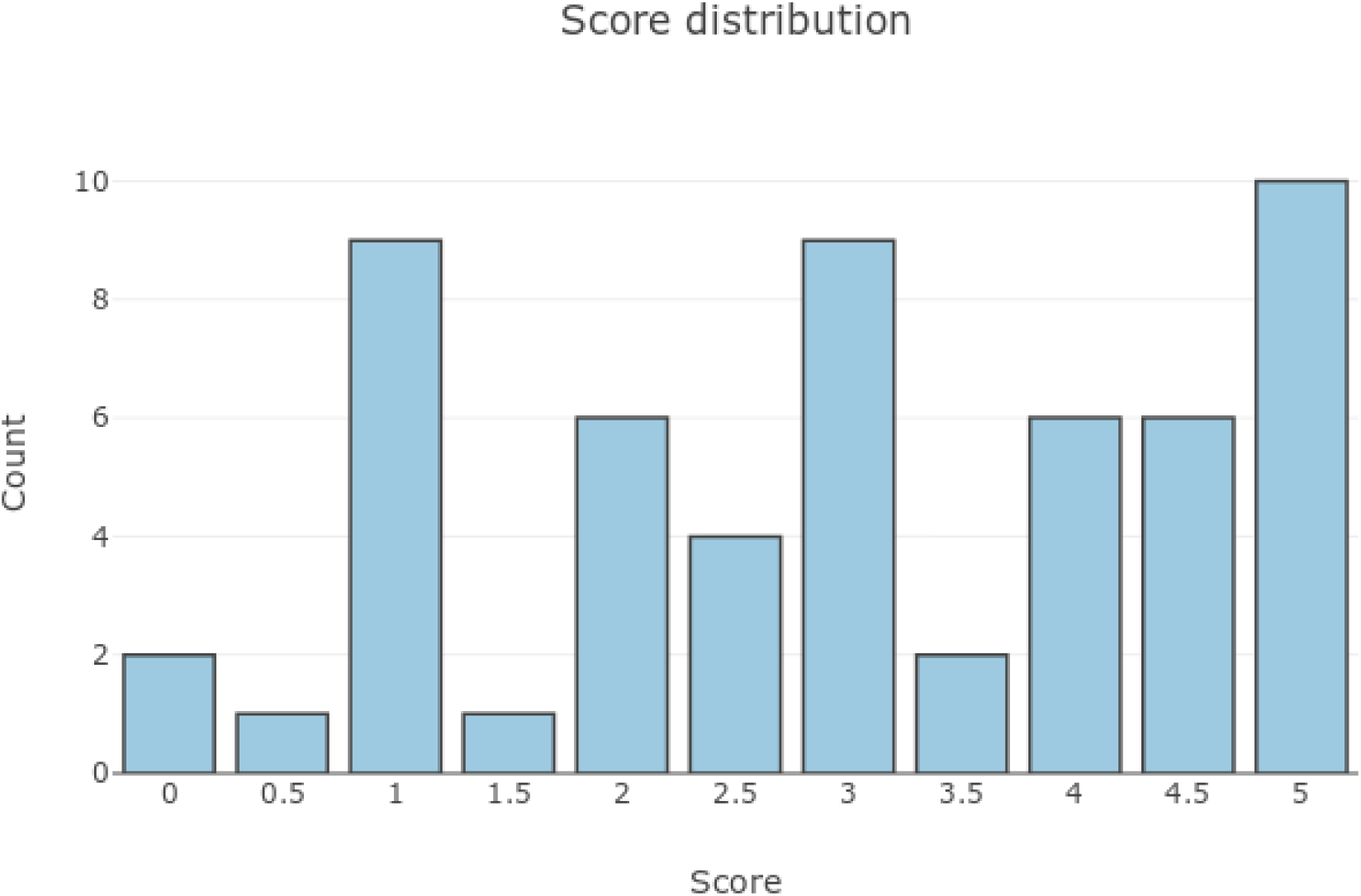
The cumulative score distribution for all evaluated resources.

(Criteria A) We found that 24 (43%) resources used an explicit standard license and 22 (39%) used custom terms, 5 (9%) had inconsistent licensing information, and 5 (9%) had no licensing information. Table 1 illustrates the count of resources by license type and the associated licensing category. Resources with custom licensing language fell into several licensing categories: 12 were restrictive, 9 permissive, and 1 private pool.

**Table 1.** Count and categories of evaluated resources’ licensing types.

| LICENSE | COUNT | CATEGORY |
| --- | --- | --- |
| custom language (restrictive) | 12 | restrictive |
| custom language (permissive) | 9 | permissive |
| CC-BY-4.0 | 8 | permissive |
| inconsistent or multiple | 5 | unknown |
| no license | 5 | copyright |
| all rights reserved | 3 | copyright |
| CC-BY-ND-3.0 | 3 | restrictive |
| CC0-1.0 | 3 | permissive |
| GPL-3.0 | 1 | copyleft |
| CC-BY-SA-3.0 | 1 | copyleft |
| CC-BY-SA-4.0 | 1 | copyleft |
| ODbL-1.0 | 1 | copyleft |
| custom language (private pool) | 1 | private pool |
| MIT | 1 | permissive |
| CC-BY-NC-4.0 | 1 | restrictive |
| public domain declaration | 1 | permissive |

(Criteria B) 32 (57%) of the resources included a license that was explicit, comprehensive, and unambiguous in scope over the data.

(Criteria C) We found that 48 (86%) of the resources passed criteria C by making all of their data reasonably accessible at an API endpoint or structured download site.

(Criteria D) We found that 20 (35%) of the data resources included clear and unambiguous licensing language that provided for unfettered reuse for all purposes

(Criteria E) 17 (30%) included clear and unambiguous language that provided for unfettered reuse for all user groups.

The majority of resources failed to receive a full star for parts A, D, or E of the rubric (Figure 3).

**Figure 3.**
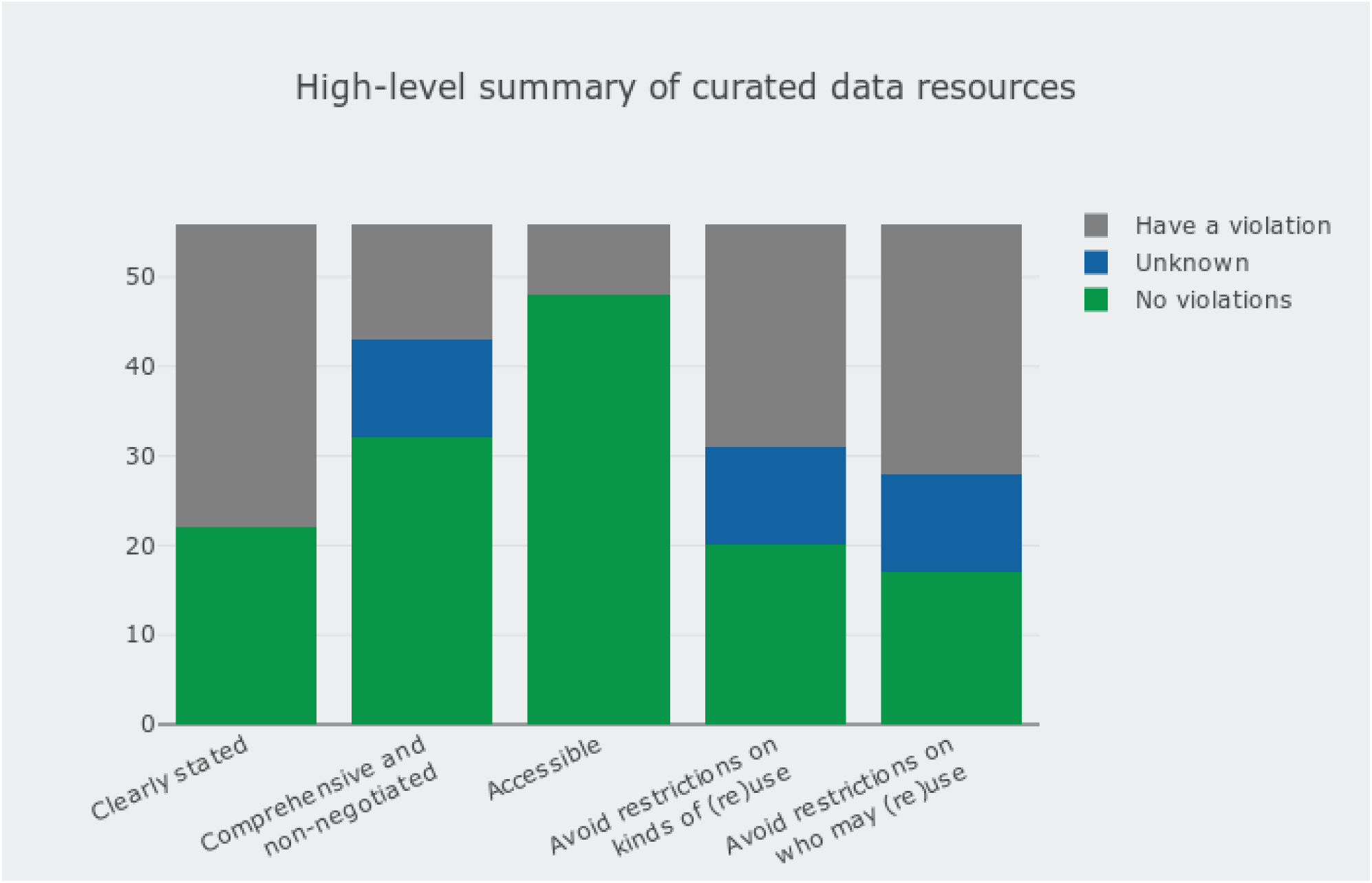
Number of data resources that passed each criteria category in the RDP rubric.

## DISCUSSION

As data users and stewards, we have encountered and been frustrated by the ways in which licensing issues hinder data reuse, integration, and redistribution. While 48 (86%) of the resources we evaluated provided easy and actionable data access, only 10 (18%) received a full 5-star rating and 32 (57%) of the resources received 3 stars or less, indicating that there were serious barriers to reuse. These findings support our experience, in that the data we need is often accessible, but cannot be reused or redistributed. Missing licensing information and the variability and potential incompatibility of license types are primary areas needing improvement. For large data integration projects that ingest data from multiple resources to derive new knowledge and provide new tools, this landscape requires costly interactions with individual organizations and institutions.

It is noteworthy that that the largest single type of licenses were custom licenses, suggesting that resource providers either felt that a standard license did not meet their needs or that they were not knowledgeable about standard licenses. Moreover, while the majority of custom licenses were restrictive, 9 were permissive, which leads us wonder if some needs and intentions are not being met by the existing set of standard permissive licenses. Although it is encouraging that the largest single license category is permissive, the total body of non-permissive license types is larger.

Our goal with the (Re)usable Data Project is to draw attention to the licensing issues that are challenging the reuse of valuable biomedical data, not to criticize any specific organization or data resource in the community. Rather, we hope RDP will encourage the community to work together to improve licensing practices in order to facilitate reusable resources for all. Reusing data *en masse* comes with numerous challenges and can be better enabled via the practices articulated in initiatives like the FAIR Data Principles and FAIR-TLC evaluation framework [15-16]. RDP’s focus on licensing issues is meant to draw attention to the pervasiveness of current practice failures and their effects.

As part of our evaluation process, we often contacted data resources with clarifying questions about their licensing information and tracked these conversations on the RDP Github repository. These exchanges led to more accurate evaluations and sparked dialogue about how resource curators could improve the clarity of their licenses. Additionally, in response to our outreach on social media, we received requests to evaluate eight additional data resources. We believe this early engagement demonstrates a community interest in enabling reuse, and a desire to contribute to open discussions about how to fix our licensing problems. Moreover, while RDP has been focused on biological and biomedical data resources, we believe the goals and problems we have raised are domain agnostic, and want to collaborate with other data communities to ensure that our rubric is relevant and applicable across disciplines.

We do not envision the RDP star rubric and evaluation data as only a tool for analyzing the past. Rather, the rubric could be used to test and guide future licensing choices. For example, it could be used by groups considering how to plan for the long-term sustainability of data resources, which may include a variety of monetization options. The RDP rubric could be applied to understand the implications of different strategies, including the potential interoperability between resources and as check on continued data reusability.

While the RDP’s star rubric and evaluation results provide a general view of the data resource licensing landscape, we believe a more in-depth approach would be valuable. We are interested, for example, in developing criteria that would define and assess more complicated interactions and compatibility characteristics between data resources. Exploring the license interaction space more deeply would require the creation of a richer internal model for our data, possibly using ontologies and leveraging the use of reasoners to aid in the task. Finally, we would like to capture and add to our analyses data resource size, connectivity, and structured funder information. These improvements could provide a more complete and holistic picture of the reusability and impact of publicly funded research data.

## Acknowledgements

We would like to thank Arvin Paranjpe, Senior Technology Development Manager, OHSU, for his thoughtful discussion and expertise on academic data licensing.

This work was support by NIH Grants OT3TR002019, U24TR002306, and R24OD011883

Seth Carbon was supported by the Director, Office of Science, Office of Basic Energy Sciences, of the U.S. Department of Energy under Contract No. DE-AC02-05CH11231”

